# Cryo-electron microscopy structure of a nucleosome-bound SWI/SNF chromatin remodeling complex

**DOI:** 10.1101/805184

**Authors:** Yan Han, Alexis A Reyes, Sara Malik, Yuan He

## Abstract

The multi-subunit chromatin remodeling complex SWI/SNF^1–3^ is highly conserved from yeast to humans and plays critical roles in various cellular processes including transcription and DNA damage repair^4, 5^. It uses the energy from ATP hydrolysis to remodel chromatin structure by sliding and evicting the histone octamer^6–10^, creating DNA regions that become accessible to other essential protein complexes. However, our mechanistic understanding of the chromatin remodeling activity is largely hindered by the lack of a high-resolution structure of any complex from this family. Here we report the first structure of SWI/SNF from the yeast *S. cerevisiae* bound to a nucleosome at near atomic resolution determined by cryo-electron microscopy (cryo-EM). In the structure, the Arp module is sandwiched between the ATPase and the Body module of the complex, with the Snf2 HSA domain connecting all modules. The HSA domain also extends into the Body and anchors at the opposite side of the complex. The Body contains an assembly scaffold composed of conserved subunits Snf12 (SMARCD/BAF60), Snf5 (SMARCB1/BAF47/ INI1) and an asymmetric dimer of Swi3 (SMARCC/BAF155/170). Another conserved subunit Swi1 (ARID1/BAF250) folds into an Armadillo (ARM) repeat domain that resides in the core of the SWI/SNF Body, acting as a molecular hub. In addition to the interaction between Snf2 and the nucleosome, we also observed interactions between the conserved Snf5 subunit and the histones at the acidic patch, which could serve as an anchor point during active DNA translocation. Our structure allows us to map and rationalize a subset of cancer-related mutations in the human SWI/SNF complex and propose a model of how SWI/SNF recognizes and remodels the +1 nucleosome to generate nucleosome-depleted regions during gene activation^11–13^.

## Main

To gain insight into the molecular mechanisms of how SWI/SNF remodels chromatin, we purified endogenous SWI/SNF from *S. cerevisiae*, assembled the SWI/SNF-nucleosome complex *in vitro* (Extended Data Fig. 1) and determined its structure using single particle cryo-EM. The complex was assembled in the presence of the non-hydrolysable ATP analog ADP-BeF_x_ and was determined to sub-nanometer resolution (Extended Data Fig. 2). We observed that the nucleosome is clamped between two regions of the SWI/SNF complex (Fig. 1, Supplementary Video 1). To improve the resolution, we also assembled the complex in the presence of ATPγS and determined its structure using cryo-EM (Extended Data Fig. 3). Since this structure shows features similar to the ADP-BeF_x_ bound complex, we combined the two data sets and performed further processing (Extended Data Fig. 4). After careful 3D classification, we obtained a reconstruction of the body of SWI/SNF to an average resolution of 4.7 Å (Fig. 1a; Extended Data Fig. 4), which we refer to as the Body module of SWI/SNF. This resolution allowed the *de novo* model building of the SWI/SNF complex (Fig. 1b).

**Fig. 1:**
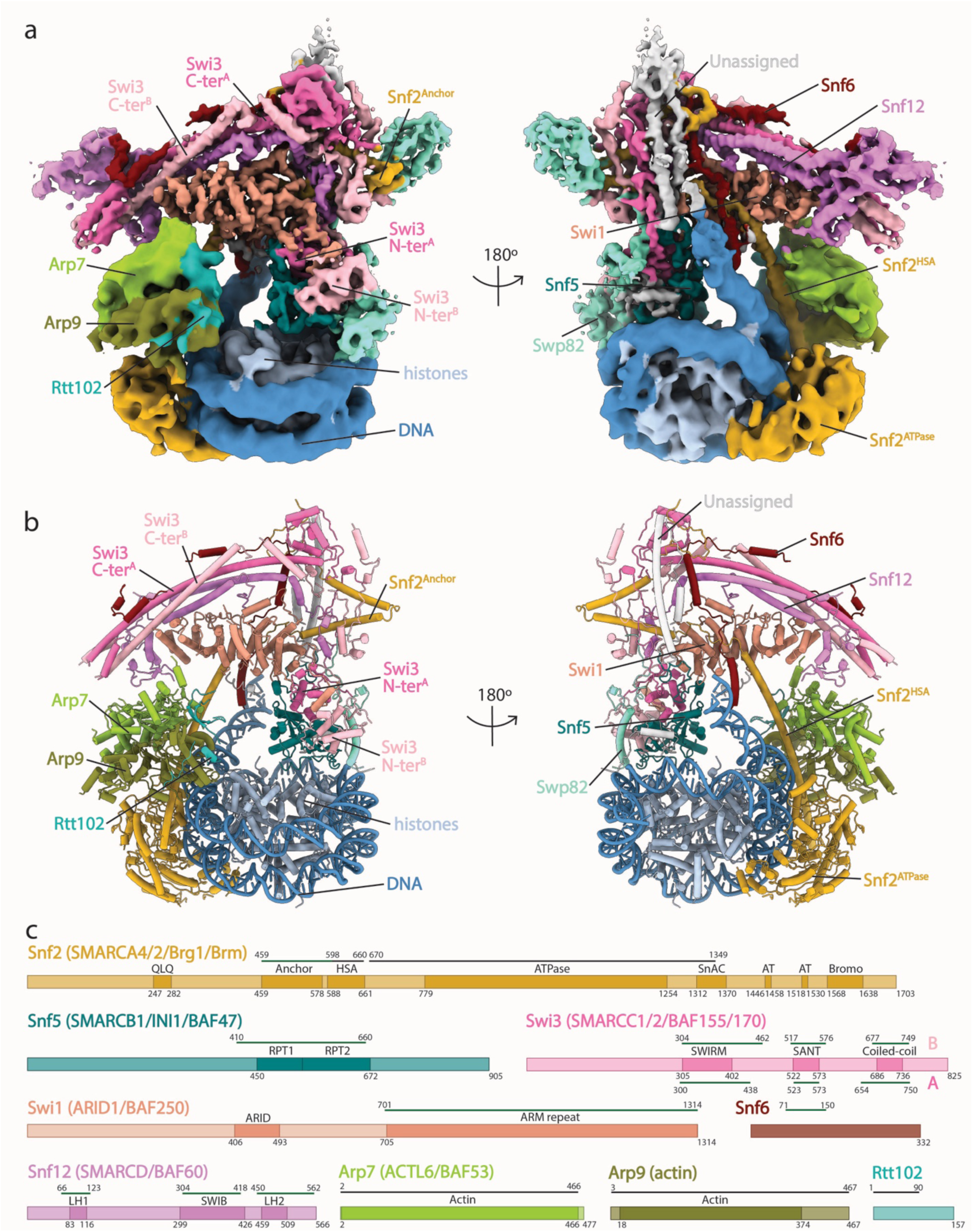
Cryo-EM structure of the SWI/SNF-nucleosome complex. **a**, Front (left) and back (right) views of the cryo-EM composite map (see Methods) of the SWI/SNF-nucleosome complex assembled in the presence of ADP-BeF_x_. **b**, Same views of structural model of the SWI/SNF-nucleosome complex as in **a**. **c**, Domain organization of all subunits that has been built in the model from **b**. Mammalian homologs are shown in parentheses. Newly built or homology regions are highlighted by green lines with residue numbers, whereas previous structures that were rigid body docked in our map are indicated by black lines. Subunits in **b** and **c** are colored as in **a**. Abbreviations: QLQ, Glutamine-Leucine-Glutamine; HSA, Helicase/SANT-associated; SnAC, Snf2 ATP coupling; AT, AT hook DNA-binding motif; RPT, Snf5 core repeat; SWIRM, a protein domain found in SWI3, RSC8 and MOIRA; SANT, SWI3, ADA2, N-CoR and TFIIIB’’ DNA-binding; ARID, AT-rich interaction domain; ARM repeat, Armadillo repeat; LH1/2, long helix 1/2; SWIB, SWI complex BAF60b.

In addition to the bound nucleosome, the SWI/SNF complex is composed of three major modules: the Body, the Arp, and the ATPase (Fig. 2a). The Snf2 ATPase domain binds the nucleosome at super helical location (SHL) 2, the same location shown in the stand-alone structures of the Snf2 ATPase-nucleosome complexes^14, 15^ as well as in SWR1^16^, Chd1^17, 18^ and SNF2h^19^, but quite different from INO80^20, 21^ (Extended Data Fig. 5). The Arp module is composed of Arp7, Arp9, Rtt102 and the HSA domain of Snf2 and is sandwiched between the Body and the ATPase modules (Fig.1a, b). This architecture has never been observed before and is quite different from other multi-subunit remodeling complexes, including INO80 and SWR1^16, 20, 21^ (Extended Data Fig. 5). The HSA of Snf2 plays an essential role in connecting the ATPase and Arp modules to the Body, extending into the Body and anchoring at the opposite side of the complex (Fig. 1a, b). We therefore named this region of Snf2 adjacent to the HSA the Anchor domain (Fig. 1c). This connection of the Arp module to the Body through a single α helix could explain the observed flexibility of the Arp and ATPase modules in the reconstruction as evidenced by lower estimated local resolution (Extended Data Figs. 2, 3). The functional relevance of this flexibility requires further investigation.

**Fig. 2:**
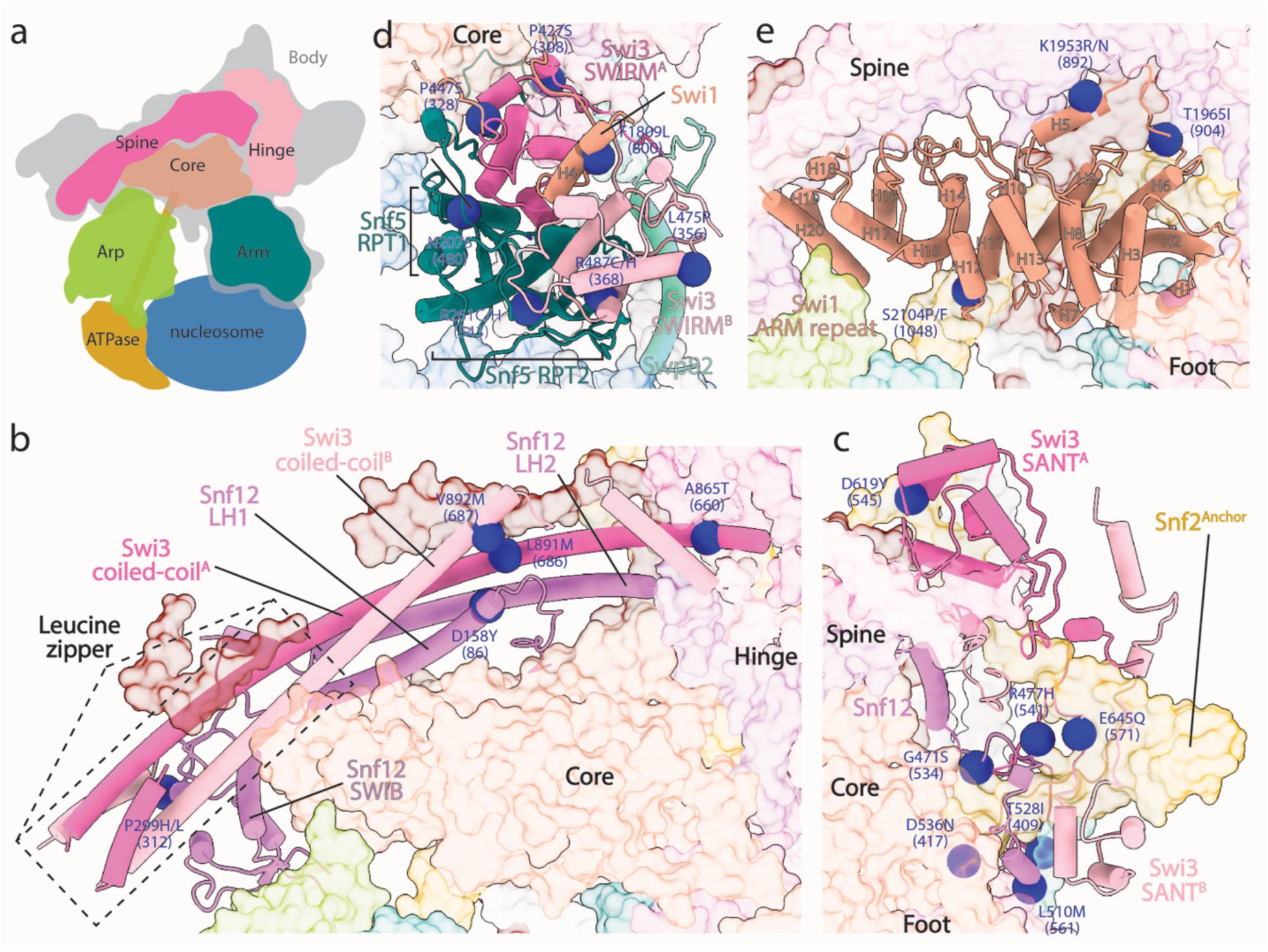
Structural organization of the Body module of SWI/SNF. **a**, A cartoon depicting the molecular architecture of the SWI/SNF-nucleosome complex. **b-e**, Close-up view of detailed interactions within the Spine (**b**), the Hinge (**c**), the Arm (**d**) and the Core (**e**) sub-modules, respectively. Blue spheres depict the locations of a subset of invariant residues harboring cancer patient mutations that occur at interfaces between these conserved subunits. Subunits are colored the same as in Fig. 1.

The 4.7 Å resolution map of the Body shows the helical nature of the SWI/SNF (Fig. 1a) and enabled us to build a structural model with the help of prior knowledge of this important complex (Figs. 1b, 2; Methods). We then mapped the crosslinking data for apo SWI/SNF^22^ onto our model of the Body module as a validation procedure (Extended Data Table 1). Out of the 35 inter-linking pairs that were mapped onto the Body model, 27 (77%) pairs have a Cα-Cα distance within 30 Å, the maximum distance that is allowed by using the crosslinker BS3^23^. We also mapped 60 pairs of intra-links, of which 55 (92%) show a Cα-Cα distance within 30 Å. These comparisons demonstrate the accuracy of our model, and also indicates that the structure of the SWI/SNF Body module does not change drastically upon engaging a nucleosome.

The conserved subunits Swi1/ARID1/BAF250, Swi3/SMARCC/BAF150/177, Snf12/ SMARCD/BAF60 and Snf5/ SMARCB1/BAF47/INI1 assemble into the body of the SWI/SNF complex (Fig. 2), consistent with these proteins forming a core module in the human SWI/SNF complexes^24^. Based on the positioning of different domains and their functions, we further defined four sub-modules of the scaffold — the Spine, the Hinge, the Arm and the Core (Fig. 2a).

The Spine is composed of Snf12 and the C-terminal regions of Swi3 (Fig. 2b). We identified two copies (named A and B) of Swi3 in our structure, consistent with previous crosslinking data showing multiple same-residue crosslinks within Swi3^22^. The most striking feature of the Spine is the four-helix bundle formed by the two long helices (LH1/2) of Snf12 and the Coiled-coil domains from two Swi3 (Fig. 2b), consistent with previous finding that the RSC homologs of Snf12 (RSC6) and Swi3 (RSC8) directly interact^25^. The Coiled-coil domain of Swi3 has clear leucine-zipper properties, containing hydrophobic amino acids separated by 7 residues in a helical region^26^. Interestingly, the crystal structure of the human dominant-negative OmoMYC homodimer^27^, a leucine-zipper containing complex, can be unambiguously fitted into the two helices belonging to Swi3 by rigid body docking (Extended Data Fig. 6a). Surprisingly, the two Coiled-coil domains of Swi3 have different lengths (Fig. 2b), showing an asymmetric folding (Extended Data Fig. 6b). We speculate that this might be due to the different interactions that the two Coiled-coils are involved in during complex assembly. BAF155/170 (SMARCC1/2), the human homologues of Swi3, have been indicated to form a dimer at the very first step of SWI/SNF complex assembly^24^. We therefore hypothesize that the two copies of Swi3 are indistinguishable at the early steps of SWI/SNF assembly, and that after engaging with other subunits, especially Snf12/SMARCD/BAF60, the symmetry is broken. Snf12 has been shown to play important roles in SWI/SNF function^28^, and our structure suggests that it may do so by interacting with Swi3 and contributing to the assembly of the complex. The unassigned density at the tip of the Spine shows clear β-sheet features and is directly connected to the SWIB domain of Snf12 (Extended Data Fig. 6c). This, together with the secondary structure prediction of Snf12, allowed us to assign this density to Snf12.

The Hinge is composed of the two SANT domains of Swi3 and the C-terminal helices of Snf12 (Fig. 2c). SANT^B^ contacts the C-terminal helices of Snf12 and is in close proximity to the Core sub-module (Fig. 2c), whereas SANT^A^ is located at the top and interacts with a C-terminal segment of Swi3^A^ (Fig. 2c). Both SANT domains contact and sandwich the Snf2 Anchor domain (Fig. 2c), playing a key role in stabilizing the ATPase within the complex.

The Arm is composed of Snf5, the N-terminal SWIRM domains of Swi3 and C-terminus of Swp82 (Fig. 2d). The Snf5 Core repeat (RPT) domains each engage one copy of the Swi3 SWIRM domain in a similar manner as in the human BAF47/BAF155 crystal structure^29^ (Fig. 2d, Extended Data Fig. 7a, b). Subtle differences in the two RPT-SWIRM interfaces (Extended Data Fig. 7c) are likely due to the α helix N-terminal to RPT1, H-N, wedging between RPT1 and SWIRM^A^ while the C-terminal region of RPT2 is packed against the opposite side of SWIRM^B^. The RPT1/SWIRM^A^ connects the Arm module to the Core module by tightly associating with Swi1 (Fig. 2d). Swp82 contains an α helix that runs along Snf5/Swi3 (Fig. 2d), likely further stabilizing the Arm module. The environments that the two molecules of Swi3 experience in both the Hinge and the Arm further establish the asymmetric architecture of this homodimer (Extended Data Fig. 6b).

Swi1/ARID1/BAF250 resides in the core region of the Body, acting as a hub to integrate all other modules (Fig. 2a, e). Therefore, we name it the Core module. It clearly folds into an Armadillo (ARM) repeat structure^30^ (Fig. 2e, Extended Data Fig. 8a). Interestingly, BAF250a, the human homolog of Swi1, was predicted to contain an ARM domain^31^, consistent with the highly conserved nature of this subunit. Compared to the β-catenin structure^32^, the Swi1 ARM repeat domain contains extra insertion sequences (Extended Data Fig. 8a), such as the one between helices H3 and H6. In addition to the neighboring repeats, this long insertion makes extensive contacts with the Snf5 and Swi3 subunits of the Arm as well as both the Spine and the Hinge. It contacts the Arm by wrapping on top of the Swi3 SWIRM^A^ domain and traveling back along the Snf5 H-N (Extended Data Fig. 8b). Interestingly, this insertion also forms an α helix H4 that contacts a surface on SWIRM^B^, whose corresponding region on SWIRM^A^ engages with Swi1 H1 and Snf5 H-N (Extended Data Fig. 7d), emphasizing the role of Swi1 in associating with the Arm module. In addition to this long insertion associating with the Arm, H1 of Swi1 contacts the SWIRM^A^ domain of Swi3, whereas H3 and H8 interact with Snf5 RPT1 (Extended Data Fig. 8b), thus connecting the Arm to the Core. The Swi1 ARM repeat domain also interacts extensively with the Spine sub-module. The entire top surface of the Swi1 ARM makes contacts with the helix bundle from the Spine, with the C-terminal helices H19 and H20 engaging the SWIB domain of Snf12 (Extended Data Fig. 8c).

The Core is also the major docking point of the Snf2 Anchor domain (Fig. 3a). H11 of Swi1 ARM interacts with an extended region of the Snf2 HSA domain that is absent from the crystal structure^33^, while H2, H6 and H9 contact the Anchor linker (Extended Data Fig. 9a). These interactions, together with the Hinge region sandwiching the Anchor helices of Snf2 (Extended Data Fig. 9b), further lock the ATPase in the complex. This observation is consistent with ARID1A being the branching subunit connecting the ATPase module with the rest of the SWI/SNF complex in humans^24^.

**Fig. 3:**
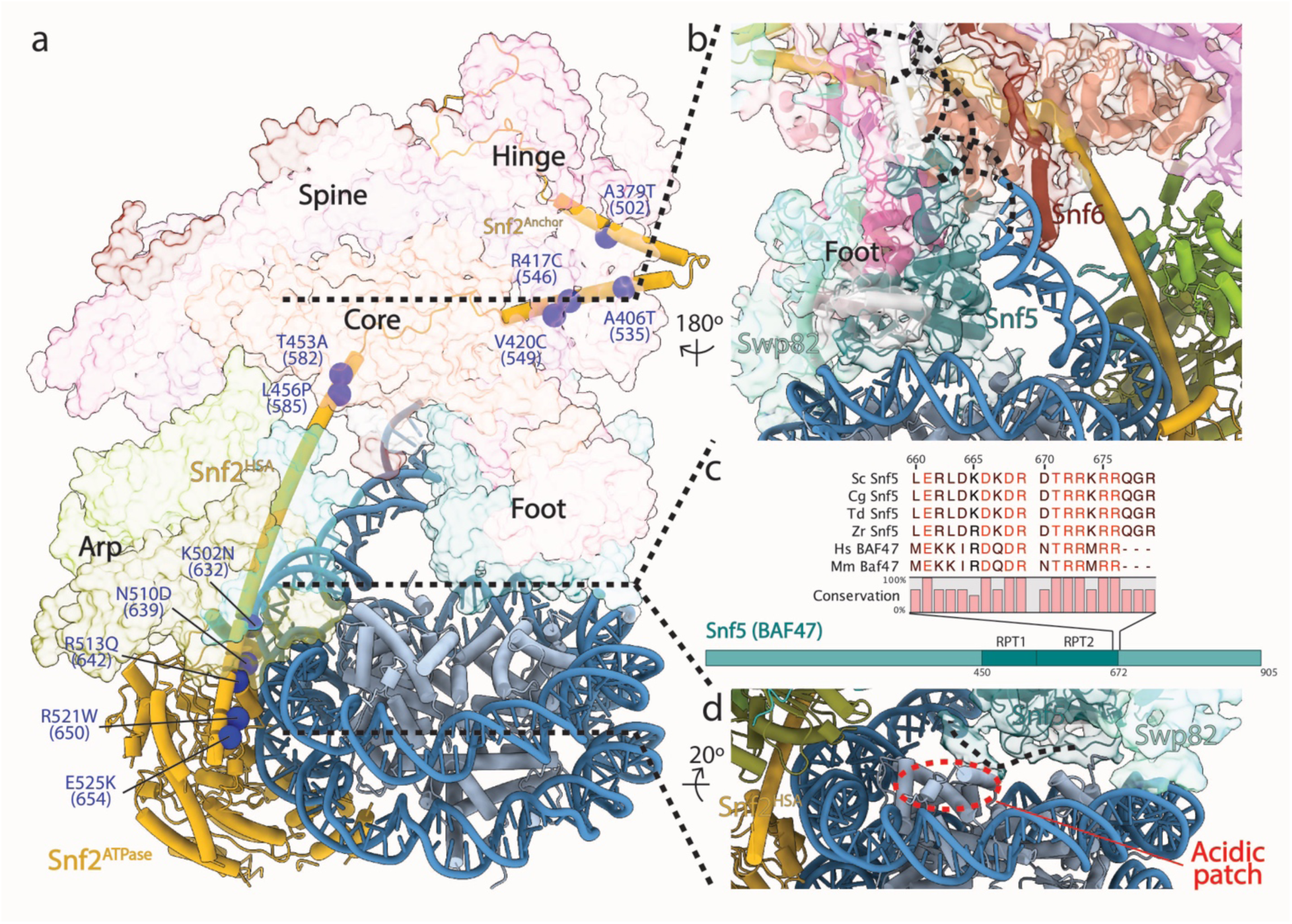
SWI/SNF-nucleosome interactions. **a**, An overview of the SWI/SNF-nucleosome complex depicting how the HSA and Anchor domains of Snf2 load the ATPase onto the nucleosome. Blue spheres indicate the positions of a subset of invariant residues harboring cancer patient mutations in Snf2/SMARCA4/BRG1 that reside between conserved SWI/SNF subunits. Dotted lines indicate regions enlarged in **b** and **d**. **b**, Close-up view showing the interaction between SWI/SNF subunits and the nucleosomal DNA. **c**, Sequence alignment of the C-terminal extension of Snf5 RPT2. Sc, *Saccharomyces cerevisiae*; Cg, *Candida glabrata*; Td, *Torulaspora delbrueckii*; Zr, *Zygosaccharomyces rouxii*; Hs, *Homo sapiens*; Mm, *Mus musculus*. Domain organization of Snf5/SMARCB1/INI1/BAF47 is also shown as in Fig. 1c. d, Close-up view showing the C-terminal extension (dotted line) of Snf5 RPT2 contacting the acidic patch of the nucleosome (dotted red circle).

The modular architecture of the SWI/SNF complex revealed by our structure agrees well with the modules revealed by previous biochemical and proteomic studies^34, 35^. The conserved SWI/SNF subunits form the structural scaffold within the complex, whereas yeast-specific subunits only occupy peripheral regions. For example, Snf6 was identified to situate at the back of the complex, spanning the Core and wrapping on top of the four-helix bundle of the Spine (Extended Data Fig. 10a). Swp82 is another yeast-specific subunit, and it is also located peripherally, making limited contacts with the rest of the complex (Extended Data Fig. 10b). Based on our sequence conservation analysis (Supplementary Figures 1-5), we have also mapped a subset of invariant residues from the human cancer mutation database^36^ onto our SWI/SNF model. Although the majority of the mutations likely compromise structure and folding, many also map to protein-protein interfaces, contributing to different types of the human disease (Fig. 2b-e, 3a).

Our structure has also enabled us to map the interactions between the SWI/SNF complex and the nucleosome. The ATPase domain of Snf2 binds the nucleosome at SHL2 in the context of the entire complex, as reported previously for the stand-alone ATPase^14, 15, 37, 38^. A series of cancer patient mutations map to the Snf2 HSA-DNA interface near SHL-6, likely diminishing the remodeling efficiency by disrupting protein-DNA interactions (Fig. 3a). The yeast-specific subunit Swp82 also contacts the nucleosomal DNA near SHL-2 (Fig. 3b), likely contributing to the remodeling activity of SWI/SNF. Although the nucleosomal DNA is not deformed as was observed for Chd1^17, 18^, there are multiple interactions between the SWI/SNF complex and the extranucleosomal DNA in our structure. First, Snf6 contacts the extranucleosomal DNA proximal to the nucleosome (Fig. 3b), in good agreement with previous site-directed DNA crosslinking experiments^39^. Second, at a lower threshold, we observed additional density for extranucleosomal DNA contacting the Body module (Extended Data Fig. 11), suggesting flexibility of this region of the DNA. However, when we prepared the SWI/SNF-nucleosome complex using a nucleosome with no overhanging DNA sequence (data not shown), we failed to observe stable complex formation, suggesting the importance of the extranucleosomal DNA in nucleosome binding to SWI/SNF. Interestingly, this extranucleosomal DNA also coincides with the possible trajectories of the N-terminal regions of both Swi1 and Snf5 (Extended Data Fig. 11), which have been shown to interact with acidic transcription activators^40–42^. This could explain how SWI/SNF is recruited by transcription activators to its target loci for chromatin remodeling, leading to an activated gene transcription.

A connecting density is observed between the histones and Snf5 C-terminus (Fig. 3c, d), consistent with the histone crosslinking experiments^22, 39^. This density likely corresponds to the highly conserved Snf5 arginine anchor motif that interacts with the acidic patch of the histone octamer (Fig. 3c, d) where a number of nucleosome regulators bind^43^, suggesting a conserved mechanism of octamer recognition. Deletion of the RPT domains in Snf5 uncouples ATP hydrolysis by Snf2 with the chromatin remodeling activity^22^. Our structure suggests an anchoring role of the Arm sub-module during active remodeling, in which Snf5 locks the histones in place as the nucleosomal DNA is being translocated, thus coupling ATP hydrolysis with chromatin remodeling (Fig. 4). In contrast, this anchoring role is primarily carried out by the Arp module in other large remodeling complexes, including INO80 and SWR1^16, 20, 21^ (Extended Data Fig. 5). It has been well documented that the natural substrate for SWI/SNF is the +1 nucleosome situated near the promoter^11–13^. Therefore, the extranucleosomal DNA at the exit side of the nucleosome in our structure corresponds to upstream promoter DNA, consistent with SWI/SNF’s function in generating the nucleosome-depleted regions during gene activation.

**Fig. 4:**
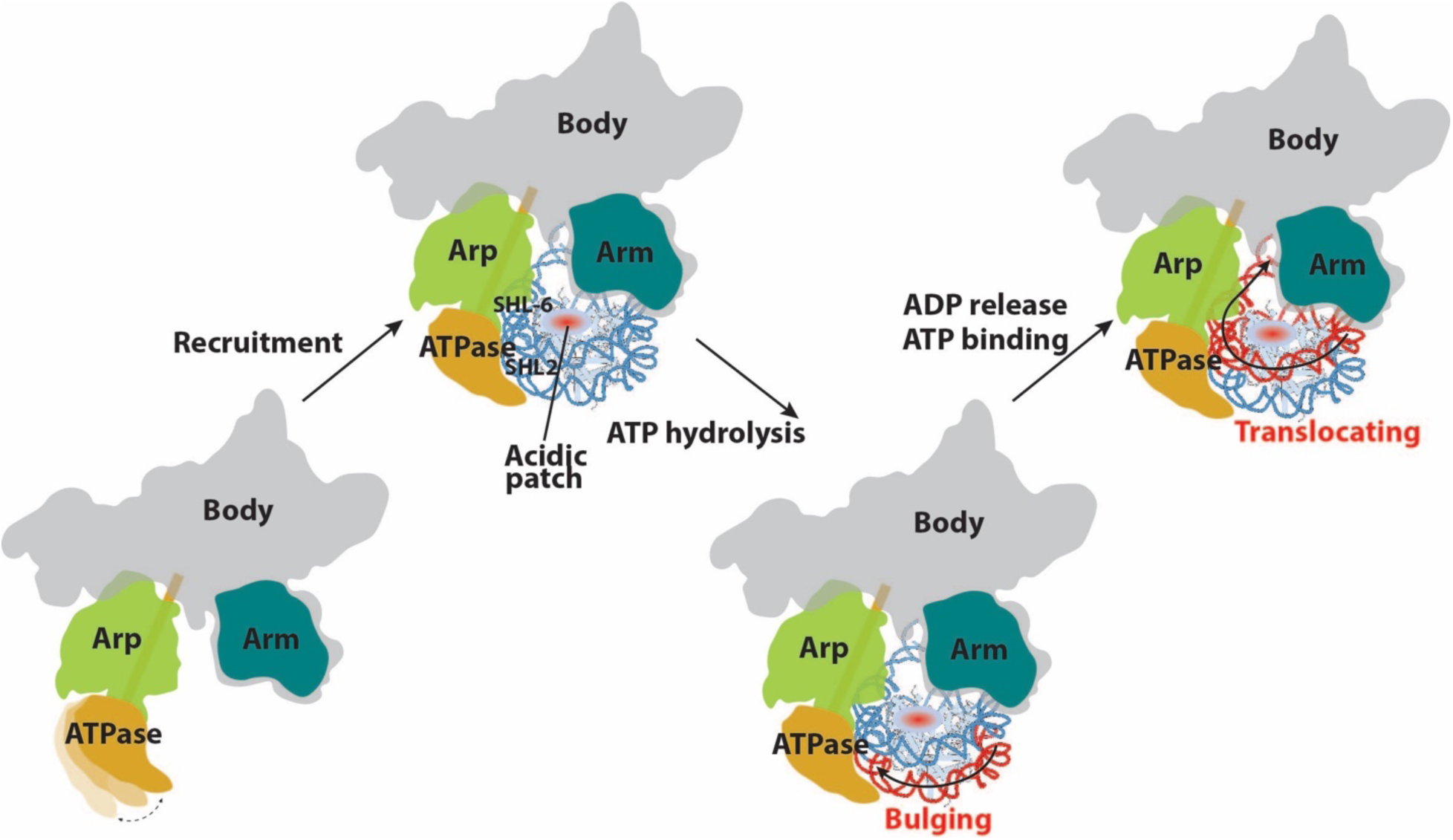
Model for chromatin remodeling activity by the SWI/SNF complex. Prior to engagement with nucleosome, the ATPase module of SWI/SNF is flexible^68^. Nucleosome binding involves the ATPase and the HSA of Snf2 recognizing SHL2 and SHL-6, respectively. The Arm of SWI/SNF interact with the nucleosome surface near the acidic patch, likely serving as an anchor during active remodeling to keep the octamer from moving. Upon ATP hydrolysis, a bulge in the nucleosomal DNA is introduced at SHL 2^15^, which is then propagated to the Exit side of the nucleosome when ADP is released and the next ATP molecule is bound, resulting in nucleosomal DNA translocation.

## Methods

### SWI/SNF purification

SWI/SNF complex was purified from a yeast strain containing a TAP tag at the C-terminus of Snf2^44^ (obtained from the High Throughput Analysis Laboratory at Northwestern University). Tandem affinity purification was performed as following. The tagged yeast strain was grown to an optical density at 600nm (OD_600_) of 4-5 in 12 liters of YPD (3% glucose). Next, cells were harvested by centrifugation and washed with 200 ml of cold TAP Extraction Buffer (40 mM Tris pH 8, 250 mM ammonium sulfate, 1mM EDTA, 10% glycerol, 0.1% Tween 20, 5 mM dithiothreitol [DTT], 2 mM phenylmethylsulfonyl fluoride [PMSF], 0.31 mg/ml benzamidine, 0.3 μg/ml leupeptin, 1.4 μg/ml pepstatin, 2 μg/ml chymostatin). Cells were resuspended in 150 ml cold TAP Extraction Buffer and lysed in a BeadBeater (Biospec Products). Cell debris was removed by centrifugation at 14,000 ×g at 4°C for 1 hr. For the first affinity step, 2 ml IgG Sepharose beads (GE Healthcare) were incubated with the lysate at 4°C overnight. The beads were next washed and resuspended in 4 ml cold TEV (tobacco etch virus) Cleavage Buffer (10 mM Tris pH8, 150 mM NaCl, 0.1% NP-40, 0.5 mM EDTA, 10% glycerol). TEV cleavage using 25 µg of TEV protease was performed at room temperature for 1 hr with gentle shaking. The TEV protease-cleaved products were collected, and the IgG beads were washed with 3 column volumes (∼6 ml total) cold Calmodulin Binding Buffer (15 mM HEPES pH7.6, 1 mM magnesium acetate, 1 mM imidazole, 2 mM CaCl_2_, 0.1% NP-40, 10% glycerol, 200 mM ammonium sulfate, 5 mM DTT, 2 mM PMSF, 0.31 mg/ml benzamidine, 0.3 μg/ml leupeptin, 1.4 μg/ml pepstatin, 2 μg/ml chymostatin). CaCl_2_ was added to the combined eluate at a final concentration of 2 mM and incubated with 0.8 ml Calmodulin Affinity Resin (Agilent Technologies) at 4°C for 2 hours. After incubation, the beads were washed with cold Calmodulin Binding Buffer and cold Calmodulin Wash Buffer (same as Calmodulin Binding Buffer, but containing 0.01% NP-40), and bound proteins were eluted with Calmodulin Elution Buffer (15 mM HEPES pH 7.6, 1 mM magnesium acetate, 1 mM imidazole, 2 mM EGTA, 10% glycerol, 0.01% NP-40, 200 mM ammonium sulfate) at room temperature. Fractions containing the SWI/SNF complex were combined and concentrated to a final concentration of ∼4 mg/ml (280 nm absorption) using a concentrator (Amicon Ultra-4 Ultracel 30K, Millipore). Concentrated protein was aliquoted, flash frozen in liquid nitrogen and stored at −80℃.

### Nucleosome reconstitution

Mono-nucleosome was reconstituted with Xenopus histones and the 601 DNA^45^ using the Mini Prep Cell (Bio-rad) as described previously^46^. The Xenopus histones were obtained from the Histone Source – the Protein Expression and Purification (PEP) Facility at Colorado State University. DNA oligonucleotides containing the 601 sequence were purchased from IDT (Integrated DNA Technology): top strand, 5’- ACCTCCCACTATTTTATGCGCCGGTATTGAACCACGCTTATGCCCAGCATCGTTA<u>ATCGATGTATATATCTGACACGTGCCTGGAGACTAGGGAGTAATCCCCTTGGCGGTTAAAACGCGGGGGACAGCGCGTACGTGCGTTTAAGCGGTGCTAGAGCTGTCTACGACCAATTGAGCGGCCTCGGCACCGGGATTCTGAT</u>-3’; bottom strand, 5’-<u>ATCAGAATCCCGGTGCCGAGGCCGCTCAATTGGTCGTAGACAGCTCTAGCACCGCTTAAACGCACGTACGCGCTGTCCCCCGCGTTTTAACCGCCAAGGGGATTACTCCCTAGTCTCCAGGCACGTGTCAGATATATACATCGAT</u>TAACGATGCTGGGCATAAGCGTGGTTCAATACCGGCGCAT-3’. The 601 sequence is underlined. The lyophilized DNA oligos were resuspended in water to a final concentration of ∼100 µM and mixed at 1:1 molar ratio. Annealing of the DNA was performed by incubating in boiling water for 5 min followed by gradually cooling to room temperature in 2 hours. The reconstituted nucleosome core particle (NCP) was concentrated to ∼6µM and annealed with a biotinylated RNA molecule (IDT, 5’-UAGUGGGAGGU-3’-biotin) to the top DNA strand at 1:1.5 (DNA to RNA) molar ratio at 45℃ for 5 min followed by gradually cooling to room temperature in 30-40 min. This resulted in a final concentration of the nucleosome core particle (NCP) at 5.52 µM. The annealed NCP was stored at 4℃.

### SWI/SNF-NCP assembly

To assemble the SWI/SNF-NCP complex, we modified our approach of reconstituting Pol I/II/III pre-initiation complexes (PIC)^47–49^ and used the NCP to replace the nucleic acid scaffold. Specifically, 0.4 µl of the biotin-RNA-annealed NCP (0.552 µM, 1/10 of the storage concentration) was first mixed with 1µl of the assembly buffer (12 mM HEPES pH 7.9, 0.12 mM EDTA, 12% glycerol, 8.25 mM MgCl_2_, 1 mM DTT, 2 mM ADP, 32mM KF, 4mM BeCl_2_ and 0.05% NP-40 [Roche]). Next, 1 µl of the concentrated SWI/SNF complex was added to this mixture and incubated at room temperature for 2 hours. Assembled complex was immobilized onto the magnetic streptavidin T1 beads (Invitrogen) which had been equilibrated with the assembly buffer plus 60 mM KCl and minus ADP-BeF_x_. Following washing of the beads two times using a wash buffer (10 mM HEPES, 10 mM Tris, pH 7.9, 5% glycerol, 5 mM MgCl_2_, 50 mM KCl, 1 mM DTT, 0.05% NP-40, 1 mM ADP, 16mM KF, 2mM BeCl_2_), the complex was eluted by incubating the beads at room temperature for 30 min with 3µl digestion buffer containing 10 mM HEPES, pH 7.9, 10 mM MgCl_2_, 50 mM KCl, 1 mM DTT, 5% glycerol, 0.05% NP-40, 1 mM ADP, 16mM KF, 2mM BeCl_2_ and 0.05 unit/µl RNase H (New England Biolabs). The SWI/SNF-NCP complex assembled in the presence of ATP*γ*S was performed essentially as described above with 1mM (2mM in the first assembly buffer) ATP*γ*S replacing ADP-BeF_x_ in the buffers.

### Electron microscopy

The assembled SWI/SNF-NCP complex was first crosslinked using 0.05% glutaraldehyde under very low illumination conditions on ice for 5 min before applied onto EM grids. Negative staining sample preparation and data collection were performed as previously described^48^. For cryo sample preparation, crosslinked complex (∼3.3 µl) was applied onto a 400 mesh Quantifoil grid containing 3.5 µm holes and 1 µm spacing (Quantifoil 3.5/1, Electron Microscopy Sciences). A thin carbon film was floated onto the grid before it was plasma cleaned for 10s at 5 W power using a Solarus plasma cleaner (Gatan) equipped with air immediately before sample deposition. The sample was allowed to absorb to the grid for 10 min at 4℃ and 100% humidity in a Vitrobot (FEI) under low illumination conditions, before blotted for 4 s at 10 force and plunge-frozen in liquid ethane. The frozen grids were stored in liquid nitrogen until imaging.

Cryo-EM data collection was performed using a JEOL 3200FS transmission electron microscope (JEOL) equipped with a K2 Summit direct electron detector (Gatan) operating at 200kV (Extended Data Table 2). Data were collected using the K2 camera in counting mode at a nominal magnification of 30,000 × (1.12 Å per pixel). Movie series with defocus values ranging from −1.5 to −4.5 µm were collected using Leginon^50^. 40-frame exposures were taken at 0.3 s per frame (12 s total), using a dose rate of 8 e^-^ per pixel per second, corresponding to a total dose of 76.5 e^-^ Å^-2^ per movie series. Four datasets with a total number of 7,769 movies on the ADP-BeF_x_ sample and four other datasets with a total number of 6,903 movies on the ATP*γ*S sample were collected.

### Image processing and three-dimensional reconstruction

Negative stain data pre-processing was performed using the Appion processing environment^51^. Particles were automatically selected from the micrographs using a difference of Gaussians (DoG) particle picker^52^. The contract transfer function (CTF) of each micrograph was estimated using CTFFind4^53^, the phases were flipped using CTFFind4, and particle stacks were extracted using a box size of 128 × 128 pixels. Two-dimensional classification was conducted using iterative multivariate statistical analysis and multi-reference alignment analysis (MSA-MRA) within the IMAGIC software^54^. Three-dimensional (3D) reconstruction of negative stained data was performed using an iterative multi-reference projection-matching approach containing libraries from the EMAN2 software package^55^. The initial 3D model was generated using cryoSPARC^56^.

Cryo-EM data was pre-processed as follows. Movie frames were aligned using MotionCor2^57^ to correct for specimen motion. Particles were automatically selected from the aligned and dose-weighted micrographs using Gautomatch (developed by Zhang K, MRC Laboratory of Molecular Biology, Cambridge, UK) with 2-fold binnning (corresponding to 2.24Å/pixel). The CTF of each micrograph and of each particle was estimated using Gctf^58^. All three-dimensional (3D) classification and refinement steps together with postprocess and local resolution estimation were performed within RELION 3.0^59^.

For the ADP-BeF_x_ dataset, 891,573 particles were automatically picked and were subjected to an initial round of 3D classification with alignment using the density obtained from negative staining as the initial reference (Extended Data Fig. 2). The “Angular sampling interval”, “Offset search range (pix)” and “Offset search step (pix)” were set to 15 degrees, 10 and 2, respectively, for the first 50 iterations. Next, these values were set back to default (7.5 degrees, 5, 1) and the 3D classification was continued until convergence. This resulted in class 3 with 198,543 particles showing sharp structural features of SWI/SNF and nucleosome. This class was subsequently refined and further classified without alignment into 5 classes with a mask around the Arp module and the nucleosome (Extended Data Fig. 2b). Class 1 with 35,214 particles from this second round of classification showed best features of the nucleosome and was chosen to proceed with 3D auto-refinement, which yielded a structure of SWI/SNF-NCP at an overall resolution of 8.96Å (Extended Data Fig. 2c). All resolutions reported herein correspond to the gold-standard Fourier shell correlation (FSC) using the 0.143 criterion^60^. The ATP*γ*S dataset with 820,117 particles was processed in a similar manner (Extended Data Fig. 3), resulting in a structure with an overall resolution of 10Å.

To focus on the Body of SWI/SNF, we combined the particles from both samples after the first round of 3D classification with a total number of 390,573 particles (Extended Data Fig. 4). Next, signal subtraction on the nucleosome and the lower half of the Arp module was performed as previously described^61^, leaving the SWI/SNF Body module and the top half of the Arp module intact. Subsequently, a 3D classification was performed with only local alignment turned on. This resulted in class 5 with 61,518 particles showing the best structural features of the Body module (Extended Data Fig. 4b). Next, we unbinned and refined the original particle stack of this class, and generated masks around the Body module, the Arp module plus the ATPase density of Snf2, and the nucleosome (Extended Data Fig. 4b). 3D multi-body refinement^62^ was then performed on this class, which drastically improved the resolution of the Body module to 4.7Å (Extended Data Fig. 4c). The core region of the Body module has a resolution close to 4.3Å (Extended Data Fig. 4b), showing densities of bulky sidechains, which enabled us to partially build the structural model of the Body module (Fig. 1b). This body map replaced its corresponding region in the ADP-BeF_x_ map to result in the composite map shown in Fig. 1a.

### Model building

To aid in model building, we performed secondary structure prediction of the SWI/SNF subunits using the Genesilico Metaserver^63^. Sequence alignment of the conserved SWI/SNF subunits were performed using CLC Sequence Viewer 7 (Supplementary Figures 1-5). To build the structural model of the SWI/SNF Body module, we first performed rigid body docking of known structures into our 4.7Å Body map. The rigid-body docking was performed in UCSF Chimera^64, 65^, which yielded good fit of the following structures: the SNF5 Repeat domains (RPTs) and Swi3 SWIRM domains from the human BAF47/BAF155 complex (PDB ID 5GJK)^29^, the SANT domain of the yeast Swi3 (PDB ID 2YUS), and the SWIB domain of mouse BAF60a (PDB ID 1UHR). The BAF47/BAF155 heterodimer and the Swi3 SANT domain can be docked in the density map at two distinct locations, indicating two copies of these domains. Indeed, chemical crosslinking combined with mass spectrometry has shown at least two copies of Swi3 in the yeast SWI/SNF complex^22^, confirming our docking experiment. Yeast Snf5 has been annotated with two SNF5 RPT domains in Pfam (http://pfam.xfam.org/protein/P18480). We observed clear density in our map that connects these two RPT domains. Next, we built homology models of these structures using Modeller^66^ and replaced the docked PDBs in the Body density map. Regions with missing or extra connecting density were then manually deleted or built in Coot^67^ based also on secondary structure predictions of these proteins.

The Snf2 Anchor domain was built manually in Coot. First, the Arp7/Arp9/Rtt102/HSA structure (PDB ID 4I6M)^33^ was rigid body docked into the full map, which helped in registering the HSA helix in the Body map. The HSA helix was then manually extended in Coot, with Y586 matching a sidechain density further confirming the register of this helix. Next, the Anchor domain was manually extended from the end of the HSA by following the connected density of the map. Again, secondary structure prediction was also used as a guide when extending the model in Coot. Bulky sidechain density at Y497, Y533 and W554 further confirmed the model.

The ARM repeat domain of Swi1 locates in the core region of the Body map with the highest local resolution, therefore enabling *de novo* model building. First, the helix density corresponding to residues 942-955 of Swi1 was chosen to model because it has the highest local resolution and that it contains a few bulky sidechain densities. Next, two α helices with poly-alanine sequence were generated in Coot, which allowed us to create a bulky residue (lysine, arginine, histidine, methionine, phenylalanine, tyrosine, and tryptophan) pattern along both directions. Subsequently, these patterns were used to search against the sequences of SWI/SNF subunits on the Sequence pattern search server (http://www-archbac.u-psud.fr/genomics/patternSearch.html), and Swi1 942-955 was one of the best hit. Further extension of this helix into connected density also matched the secondary structures of Swi1. Then, the remaining regions of the Swi1 ARM repeat domain were manually built into the density in Coot based on secondary structure prediction as well as bulky sidechain densities wherever possible. The overall architecture of the ARM repeat domain of Swi1 also matches that of an Armadillo repeat containing protein β-catenin^32^ (Extended Data Fig. 8a), confirming our model of Swi1.

The positioning of the SWIB domain of Snf12 aided us in building the remaining of this protein into the density. First, at the Spine tip, where the SWIB was docked, there is β-sheet like density (Extended Data Fig. 6c). This agrees with the secondary structure prediction of Snf12, which shows β-strands right N-terminus of the SWIB domain.

Although the resolution of this region is low, we are confident about its identity. Four long helices belonging to the Spine module directly connect to this region, two of which extending into the Snf12 densities. Therefore, we assigned these two helices to Snf12. This agreed well with the secondary structure prediction of Snf12, which shows that Snf12 contains two long helices. Next, we performed protein sequence pattern search based on the bulky sidechain densities. Based on the search results, we manually built the two helices of Snf12 in Coot.

Based on secondary structure prediction, we reasoned that the other two long helices belong to Swi3 C-terminus. This is backed up by the finding that the C-terminus of Swi3 contains a coiled-coil leucine zipper motif^26^ and there are two copies of Swi3 in SWI/SNF. To facilitate the registering of the sequence in these long helices, we fitted the crystal structure of human OmoMYC homodimer (PDB ID 5I4Z)^27^ into the density and obtained a good fit (Extended Data Fig. 6a). Based on this fitting, we mapped the hydrophobic residues from Swi3 as indicated before^26^ and manually built the two helices in Coot. The rest of Swi3 density cannot be confidently modeled due to lower resolution and missing density, therefore are modeled with poly-alanine.

Snf6 was also manually built in Coot based on secondary structure prediction, bulky sidechain density and prior knowledge based on chemical crosslinking and mass spectrometry data^22^ and site-directed DNA crosslinking experiments^39^. We cannot confidently model in Swp82, however we were able to assign densities to this yeast specific subunit based on crosslinking experiments^22^ and mapping by deletions and EM^68^. The N-terminal region of Swp82 forms a RSC7 homology domain, therefore we speculate that it occupies the globular density near the Hinge; C-terminal region crosslinks to both Snf5 and Swi3, therefore it was assigned to the density by Snf5 and Swi3. There are also several unassigned densities on the solvent exposed surface of the complex. We did not identify Taf14 and Snf11 in the map (Extended Data Table 3).

The molecular model of SWI/SNF Body module was then refined using Namdinator^69^ (Extended Data Table 2). To obtain the model for the full complex, we rigid-body fitted the Body, the Arp module (PDB ID 4I6M)^33^ and the ATPase-nucleosome bound with ADP-BeF_x_ (PDB ID 5Z3V)^15^ into the map of the full complexes in Coot. Then, the HSA helix was connected manually, and the DNA sequence was modified to match our sequence. The extra DNA was manually extended by 10bp using B form DNA in Coot. The figures were prepared using UCSF Chimera and ChimeraX^70^. Cα-Cα distances from crosslinked lysine pairs^22^ were measured in UCSF Chimera. For crosslinks involving the two molecules of Swi3, we picked the combination that gave the shortest distance as the measurement (Extended Data Table 1).

## Data availability

Cryo-EM density maps have been deposited in the Electron Microscopy Data Bank (EMDB) under accession numbers EMD-XXXX (ADP-BeF_x_), EMD-XXXX (ATP*γ*S), EMD-XXXX (body). Model coordinates have been deposited in the Protein Data Bank (PDB) under accession numbers XXXX (ADP-BeF_x_), XXXX (body).

## Supporting information

Extend Data Figures

## Acknowledgement

We thank Dr. Jonathan Remis for assistance with microscope operation and data collection and Jason Pattie for computer support. We are grateful to Amy Rosenzweig, Ishwar Radhakrishnan and Susan Fishbain for helpful discussion and comments on the manuscript. We also thank the staff at the Structural Biology Facility (SBF) of Northwestern University for technical support. This work was supported by a Cornew Innovation Award from the Chemistry of Life Processes Institute at Northwestern University (to Y He), a Catalyst Award by the Chicago Biomedical Consortium with support from the Searle Funds at The Chicago Community Trust (to Y He), an Institutional Research Grant from the American Cancer Society (IRG-15-173-21 to Y He), an H Foundation Core Facility Pilot Project Award (to Y He). Y He is supported by P01 CA092584 and U54CA193419 from NIH/NCI. Y Han is a recipient of the Chicago Biomedical Consortium Postdoctoral Research Grant.

## Author contributions

Y Han and Y He conceived the project. Y Han performed most of the experiments and collected and analyzed cryo-EM data with Y He. AA Reyes and S Malik contributed to protein purification. Y Han built the models with help from Y He. Y Han and Y He wrote the manuscript, with input from all other authors.

## References

1. Côté, J., Quinn, J., Workman, J. L. & Peterson, C. L. Stimulation of GAL4 derivative binding to nucleosomal DNA by the yeast SWI/SNF complex. Science 265, 53–60 (1994).

2. Cairns, B. R., Kim, Y. J., Sayre, M. H., Laurent, B. C. & Kornberg, R. D. A multisubunit complex containing the SWI1/ADR6, SWI2/SNF2, SWI3, SNF5, and SNF6 gene products isolated from yeast. Proc. Natl. Acad. Sci. U.S.A. 91, 1950– 1954 (1994).

3. Peterson, C. L., Dingwall, A. & Scott, M. P. Five SWI/SNF gene products are components of a large multisubunit complex required for transcriptional enhancement. Proc. Natl. Acad. Sci. U.S.A. 91, 2905–2908 (1994).

4. Bartholomew, B. Regulating the chromatin landscape: structural and mechanistic perspectives. Annu. Rev. Biochem. 83, 671–696 (2014).

5. Clapier, C. R. & Cairns, B. R. The biology of chromatin remodeling complexes. Annu. Rev. Biochem. 78, 273–304 (2009).

6. Cairns, B. R. Chromatin remodeling machines: similar motors, ulterior motives. Trends in Biochemical Sciences 23, 20–25 (1998).

7. Kingston, R. E., Bunker, C. A. & Imbalzano, A. N. Repression and activation by multiprotein complexes that alter chromatin structure. Genes & Development 10, 905–920 (1996).

8. Peterson, C. L. & Tamkun, J. W. The SWI-SNF complex: a chromatin remodeling machine? Trends in Biochemical Sciences 20, 143–146 (1995).

9. Vignali, M., Hassan, A. H., Neely, K. E. & Workman, J. L. ATP-dependent chromatin-remodeling complexes. Molecular and Cellular Biology 20, 1899–1910 (2000).

10. Narlikar, G. J., Sundaramoorthy, R. & Owen-Hughes, T. Mechanisms and functions of ATP-dependent chromatin-remodeling enzymes. Cell 154, 490–503 (2013).

11. Rando, O. J. & Winston, F. Chromatin and transcription in yeast. Genetics 190, 351–387 (2012).

12. Rawal, Y. et al. SWI/SNF and RSC cooperate to reposition and evict promoter nucleosomes at highly expressed genes in yeast. Genes & Development 32, 695– 710 (2018).

13. Kubik, S. et al. Opposing chromatin remodelers control transcription initiation frequency and start site selection. Nature Structural & Molecular Biology 26, 744– 754 (2019).

14. Liu, X., Li, M., Xia, X., Li, X. & Chen, Z. Mechanism of chromatin remodelling revealed by the Snf2-nucleosome structure. Nature 544, 440–445 (2017).

15. Li, M. et al. Mechanism of DNA translocation underlying chromatin remodelling by Snf2. Nature 567, 409–413 (2019).

16. Willhoft, O. et al. Structure and dynamics of the yeast SWR1-nucleosome complex. Science 362, eaat7716 (2018).

17. Farnung, L., Vos, S. M., Wigge, C. & Cramer, P. Nucleosome-Chd1 structure and implications for chromatin remodelling. Nature 550, 539–542 (2017).

18. Sundaramoorthy, R. et al. Structural reorganization of the chromatin remodeling enzyme Chd1 upon engagement with nucleosomes. Elife 6, 1405 (2017).

19. Armache, J.-P. et al. Cryo-EM structures of remodeler-nucleosome intermediates suggest allosteric control through the nucleosome. Elife 8, 213 (2019).

20. Eustermann, S. et al. Structural basis for ATP-dependent chromatin remodelling by the INO80 complex. Nature 556, 386–390 (2018).

21. Ayala, R. et al. Structure and regulation of the human INO80-nucleosome complex. Nature 556, 391–395 (2018).

22. Sen, P. et al. Loss of Snf5 Induces Formation of an Aberrant SWI/SNF Complex. Cell Rep 18, 2135–2147 (2017).

23. Merkley, E. D. et al. Distance restraints from crosslinking mass spectrometry: mining a molecular dynamics simulation database to evaluate lysine-lysine distances. Protein Sci. 23, 747–759 (2014).

24. Mashtalir, N. et al. Modular Organization and Assembly of SWI/SNF Family Chromatin Remodeling Complexes. Cell 175, 1272–1288.e20 (2018).

25. Treich, I., Ho, L. & Carlson, M. Direct interaction between Rsc6 and Rsc8/Swh3,two proteins that are conserved in SWI/SNF-related complexes. Nucleic Acids Research 26, 3739–3745 (1998).

26. Wang, W. et al. Diversity and specialization of mammalian SWI/SNF complexes. Genes & Development 10, 2117–2130 (1996).

27. Jung, L. A. et al. OmoMYC blunts promoter invasion by oncogenic MYC to inhibit gene expression characteristic of MYC-dependent tumors. Oncogene 36, 1911– 1924 (2017).

28. Cairns, B. R., Levinson, R. S., Yamamoto, K. R. & Kornberg, R. D. Essential role of Swp73p in the function of yeast Swi/Snf complex. Genes & Development 10, 2131–2144 (1996).

29. Yan, L., Xie, S., Du, Y. & Qian, C. Structural Insights into BAF47 and BAF155 Complex Formation. Journal of Molecular Biology 429, 1650–1660 (2017).

30. Peifer, M., Berg, S. & Reynolds, A. B. A repeating amino acid motif shared by proteins with diverse cellular roles. Cell 76, 789–791 (1994).

31. Sandhya, S., Maulik, A., Giri, M. & Singh, M. Domain architecture of BAF250a reveals the ARID and ARM-repeat domains with implication in function and assembly of the BAF remodeling complex. PLoS ONE 13, e0205267 (2018).

32. Huber, A. H., Nelson, W. J. & Weis, W. I. Three-dimensional structure of the armadillo repeat region of beta-catenin. Cell 90, 871–882 (1997).

33. Schubert, H. L. et al. Structure of an actin-related subcomplex of the SWI/SNF chromatin remodeler. PNAS 110, 3345–3350 (2013).

34. Yang, X., Zaurin, R., Beato, M. & Peterson, C. L. Swi3p controls SWI/SNF assembly and ATP-dependent H2A-H2B displacement. Nat Struct Mol Biol 14, 540–547 (2007).

35. Dutta, A. et al. Composition and Function of Mutant Swi/Snf Complexes. Cell Rep 18, 2124–2134 (2017).

36. Tate, J. G. et al. COSMIC: the Catalogue Of Somatic Mutations In Cancer. Nucleic Acids Research 47, D941–D947 (2019).

37. Zofall, M., Persinger, J., Kassabov, S. R. & Bartholomew, B. Chromatin remodeling by ISW2 and SWI/SNF requires DNA translocation inside the nucleosome. Nat Struct Mol Biol 13, 339–346 (2006).

38. Saha, A., Wittmeyer, J. & Cairns, B. R. Chromatin remodeling through directional DNA translocation from an internal nucleosomal site. Nat Struct Mol Biol 12, 747– 755 (2005).

39. Dechassa, M. L. et al. Architecture of the SWI/SNF-nucleosome complex. Molecular and Cellular Biology 28, 6010–6021 (2008).

40. Neely, K. E., Hassan, A. H., Brown, C. E., Howe, L. & Workman, J. L. Transcription activator interactions with multiple SWI/SNF subunits. Molecular and Cellular Biology 22, 1615–1625 (2002).

41. Prochasson, P., Neely, K. E., Hassan, A. H., Li, B. & Workman, J. L. Targeting activity is required for SWI/SNF function in vivo and is accomplished through two partially redundant activator-interaction domains. MOLCEL 12, 983–990 (2003).

42. Mechanism of transcription factor recruitment by acidic activators. J. Biol. Chem. 280, 21779–21784 (2005).

43. McGinty, R. K. & Tan, S. Recognition of the nucleosome by chromatin factors and enzymes. Curr. Opin. Struct. Biol. 37, 54–61 (2016).

44. Ghaemmaghami, S. et al. Global analysis of protein expression in yeast. Nature 425, 737–741 (2003).

45. Lowary, P. T. & Widom, J. New DNA sequence rules for high affinity binding to histone octamer and sequence-directed nucleosome positioning. Journal of Molecular Biology 276, 19–42 (1998).

46. Dyer, P. N. et al. Reconstitution of nucleosome core particles from recombinant histones and DNA. Meth. Enzymol. 375, 23–44 (2004).

47. He, Y. et al. Near-atomic resolution visualization of human transcription promoter opening. Nature 533, 359–365 (2016).

48. Knutson, B. A. Structural mechanism of ATP-independent transcription initiation by RNA polymerase I. Elife 6, 753 (2017).

49. Han, Y., Yan, C., Fishbain, S., Ivanov, I. & He, Y. Structural visualization of RNA polymerase III transcription machineries. Cell Discov 4, 40 (2018).

50. Suloway, C. et al. Automated molecular microscopy: the new Leginon system. Journal of Structural Biology 151, 41–60 (2005).

51. Lander, G. C. et al. Appion: an integrated, database-driven pipeline to facilitate EM image processing. Journal of Structural Biology 166, 95–102 (2009).

52. Voss, N. R., Yoshioka, C. K., Radermacher, M., Potter, C. S. & Carragher, B. DoG Picker and TiltPicker: software tools to facilitate particle selection in single particle electron microscopy. Journal of Structural Biology 166, 205–213 (2009).

53. Rohou, A. & Grigorieff, N. CTFFIND4: Fast and accurate defocus estimation from electron micrographs. Journal of Structural Biology 192, 216–221 (2015).

54. van Heel, M., Harauz, G., Orlova, E. V., Schmidt, R. & Schatz, M. A new generation of the IMAGIC image processing system. Journal of Structural Biology 116, 17–24 (1996).

55. Tang, G. et al. EMAN2: an extensible image processing suite for electron microscopy. Journal of Structural Biology 157, 38–46 (2007).

56. Punjani, A., Rubinstein, J. L., Fleet, D. J. & Brubaker, M. A. cryoSPARC: algorithms for rapid unsupervised cryo-EM structure determination. Nature Methods 14, 290–296 (2017).

57. Zheng, S. Q. et al. MotionCor2: anisotropic correction of beam-induced motion for improved cryo-electron microscopy. Nature Methods 14, 331–332 (2017).

58. Zhang, K. Gctf: Real-time CTF determination and correction. Journal of Structural Biology 193, 1–12 (2016).

59. Kimanius, D., Forsberg, B. O., Scheres, S. H. & Lindahl, E. Accelerated cryo-EM structure determination with parallelisation using GPUs in RELION-2. Elife 5, e18722 (2016).

60. Henderson, R. et al. Outcome of the first electron microscopy validation task force meeting. in 20, 205–214 (2012).

61. Bai, X.-C., Rajendra, E., Yang, G., Shi, Y. & Scheres, S. H. W. Sampling the conformational space of the catalytic subunit of human γ-secretase. Elife 4, 1485 (2015).

62. Nakane, T., Kimanius, D., Lindahl, E. & Scheres, S. H. Characterisation of molecular motions in cryo-EM single-particle data by multi-body refinement in RELION. Elife 7, 1485 (2018).

63. Kurowski, M. A. & Bujnicki, J. M. GeneSilico protein structure prediction meta-server. Nucleic Acids Research 31, 3305–3307 (2003).

64. Pettersen, E. F. et al. UCSF Chimera--a visualization system for exploratory research and analysis. J Comput Chem 25, 1605–1612 (2004).

65. Goddard, T. D., Huang, C. C. & Ferrin, T. E. Visualizing density maps with UCSF Chimera. Journal of Structural Biology 157, 281–287 (2007).

66. Webb, B. & Sali, A. Comparative Protein Structure Modeling Using MODELLER. Curr Protoc Bioinformatics 54, 5.6.1–5.6.37 (2016).

67. Emsley, P., Lohkamp, B., Scott, W. G. & Cowtan, K. Features and development of Coot. Acta Crystallogr. D Biol. Crystallogr. 66, 486–501 (2010).

68. Zhang, Z. et al. Architecture of SWI/SNF chromatin remodeling complex. Protein & Cell 15, 1–5 (2018).

69. Kidmose, R. T. et al. Namdinator - automatic molecular dynamics flexible fitting of structural models into cryo-EM and crystallography experimental maps. IUCrJ 6, 526–531 (2019).

70. Goddard, T. D. et al. UCSF ChimeraX: Meeting modern challenges in visualization and analysis. Protein Sci. 27, 14–25 (2018).

