## Supplementary material for "Cryo-electron microscopy structure of a nucleosome-bound SWI/SNF chromatin remodeling complex": Extend Data Figures

**Extended Data Figures.**

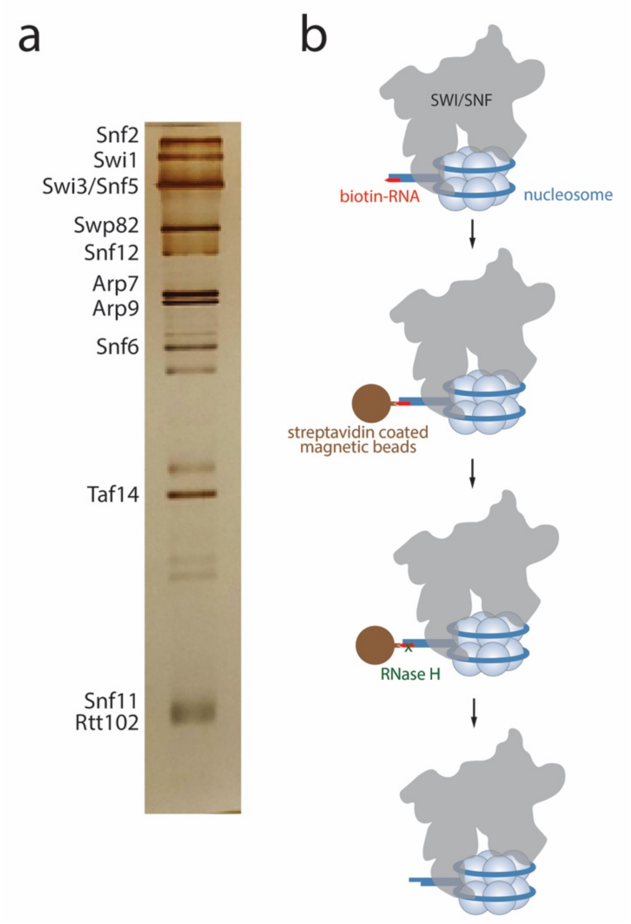


**Extended Data Fig. 1 Purification and assembly of the SWI/SNF-nucleosome complex. a**, A silver stained protein SDS-PAGE gel showing the TAP-purified SWI/SNF complex from the yeast strain bearing a TAP tag at the C-terminus of Snf2. SWI/SNF subunits are labeled based on molecular weight. **b**, Schematic showing the assembly and purification protocol of the SWI/SNF-nucleosome complex prior to single particle cryo-EM analysis. SWI/SNF is first incubated with reconstituted nucleosome. The nucleosomal DNA contains a single-stranded overhang that is annealed to a biotinylated RNA molecule. Next, the assembled complex is immobilized onto streptavidin coated magnetic beads. Following washes, the complex is eluted using RNase H digestion. The eluted complex is then crosslinked and deposited onto EM grid for vitrification.

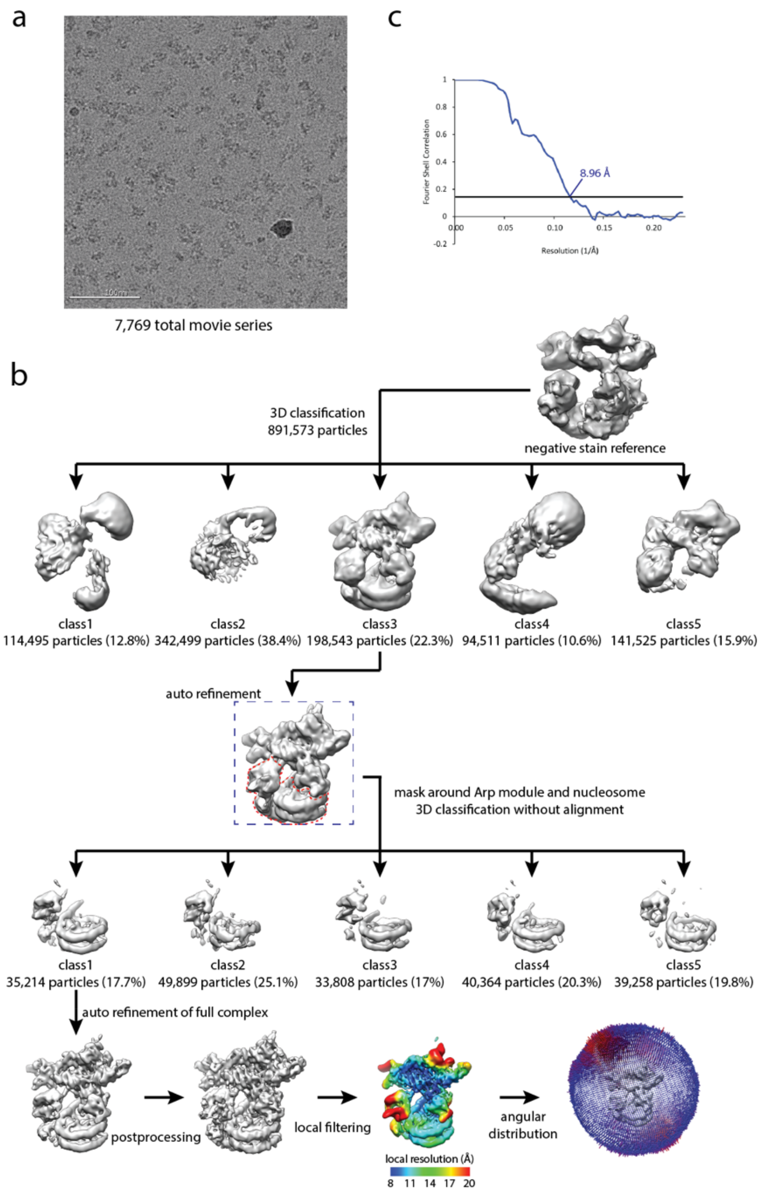


**Extended Data Fig. 2 Data processing scheme of the ADP-BeF$_x$ sample. a**, A representative raw micrograph of the SWI/SNF-nucleosome complex assembled in the presence of ADP-BeF$_x$. **b**, Flow chart of the cryo-EM data processing procedure. The particle stack of class 3 (198,543 particles; blue dashed box) after the first sorting was chosen for the combined processing (Extended Data Fig. 4). **c**, Fourier Shell Correlation curve of the complex showing a final average resolution of 8.96Å (FSC=0.143).

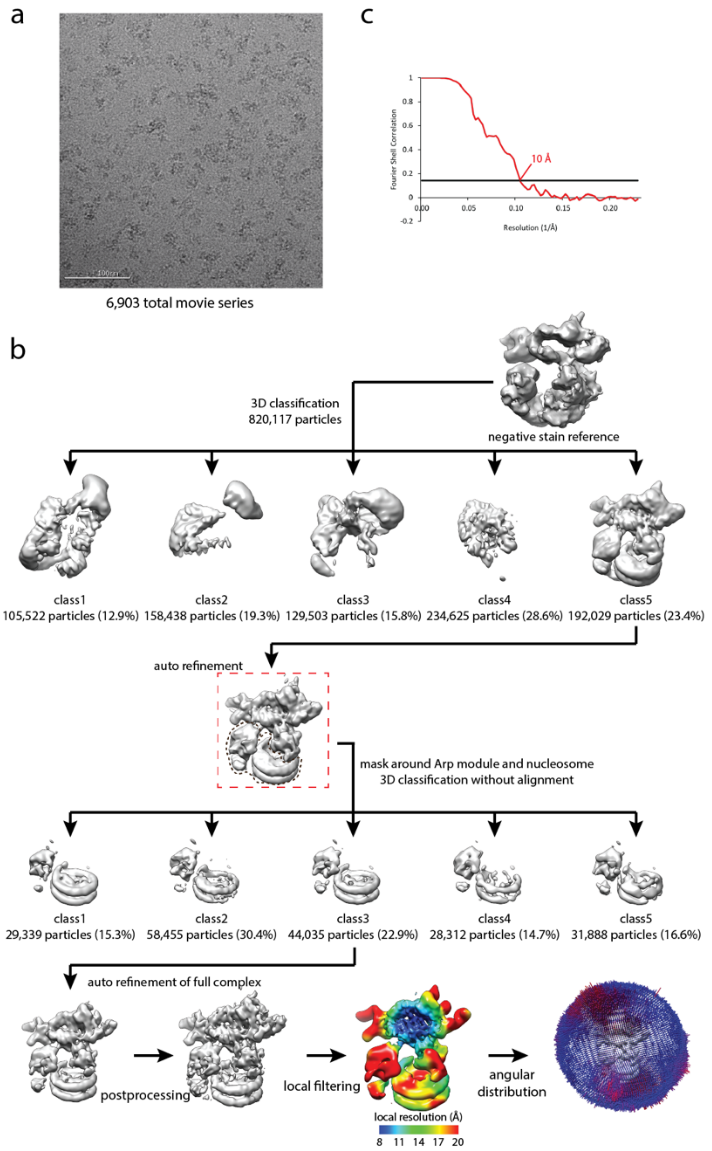


**Extended Data Fig. 3 Data processing scheme of the ATPγS sample. a**, A representative raw micrograph of the SWI/SNF-nucleosome complex assembled in the presence of ATPγS. **b**, Flow chart of the cryo-EM data processing procedure. The particle stack of class 5 (192,029 particles; red dashed box) after the first sorting was chosen for the combined processing (Extended Data Fig. 4). **c**, Fourier Shell Correlation curve of the complex showing a final average resolution of 10Å (FSC=0.143).

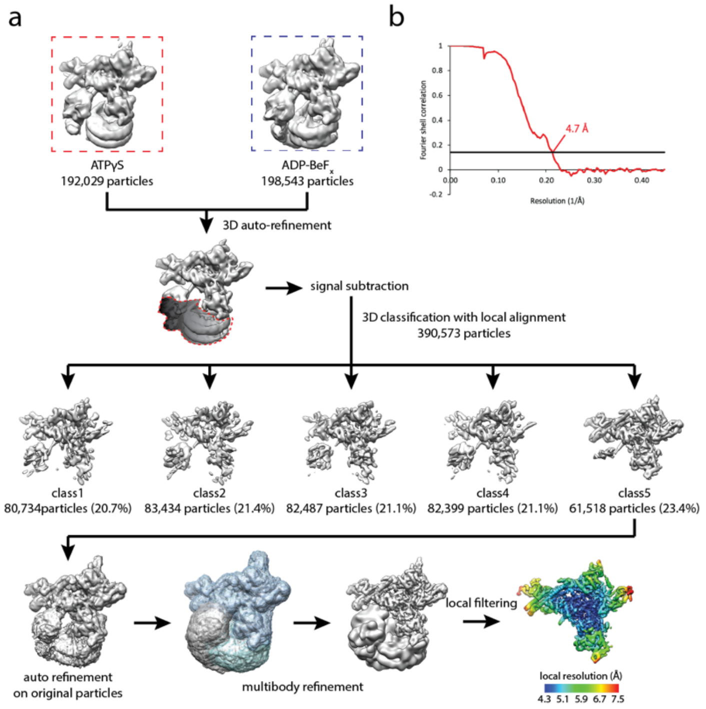


**Extended Data Fig. 4 Data processing scheme of the combined dataset. a**, Flow chart of the data processing procedure by combining ADP-$BeF_x$ (Extended Data Fig. 2) and ATPγS (Extended Data Fig. 3) datasets.. **b**, Fourier Shell Correlation curve of the Body module showing a final average resolution of 4.7Å (FSC=0.143).

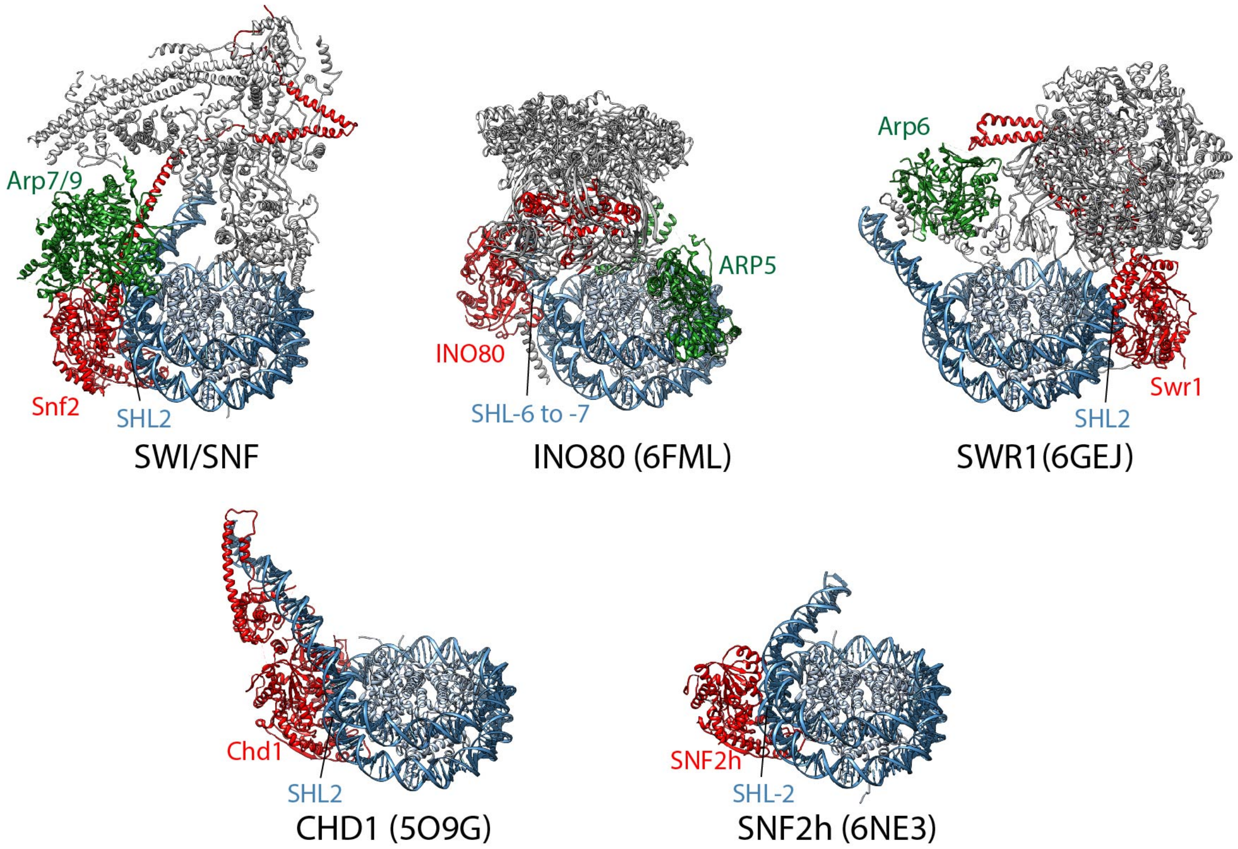


**Extended Data Fig. 5 Comparing the structures of chromatin remodelers from different families.** All but the INO80 complex have their ATPase module binding to SHL2/-2. INO80 engages the nucleosomal DNA at SHL -6 to -7. SWI/SNF is different from the INO80/SWR1 family remodelers in that its Arp module (Arp7/9) is sandwiched between the Body and the ATPase modules. The Snf2 ATPase module is connected through the long HSA domain to the rest of the complex, whereas the ATPases INO80 and Swr1 directly contact the main body of the corresponding complexes. All remodelers are aligned based on histone proteins. The ATPase in each complex is colored red, whereas Arp proteins are colored green. PDB codes of the other chromatin remodelers are shown in parentheses.

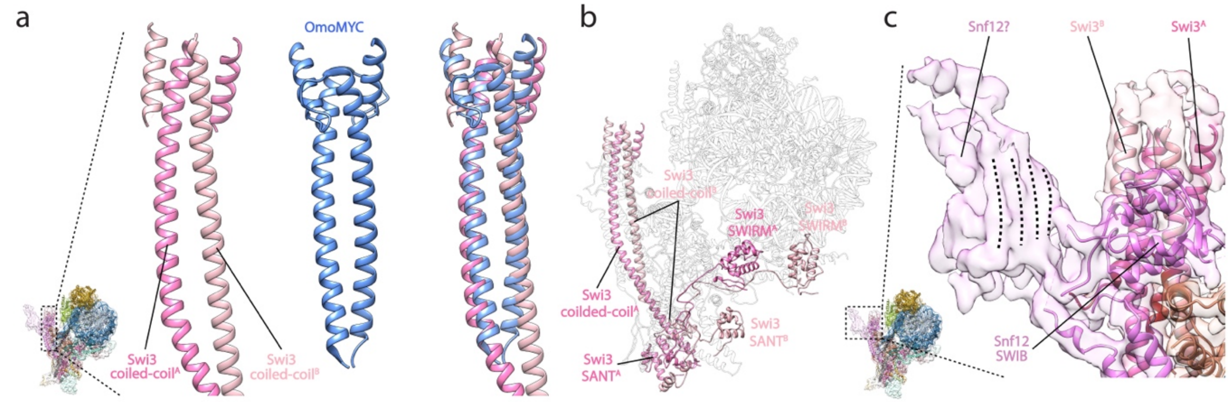


**Extended Data Fig. 6 Structural features of the Spine sub-module of the SWI/SNF complex. a**, The Swi3 Coiled-coil dimer (left) resembles the structure of the dominant-negative allele of MYC (OmoMYC, shown in blue in the middle panel; PDB ID 5I4Z). The OmoMYC structure was rigid body docked in the Spine density corresponding to the Swi3 Coiled-coil and then compared with the Swi3 Coiled-coil. **b**, Swi3 forms an asymmetric dimer in the SWI/SNF complex. **c**, The density at the tip of the Spine shows features of β-sheet and is therefore assigned to Snf12 based on closed proximity to Snf12 SWIB domain and secondary structure prediction (Supplementary Figures).

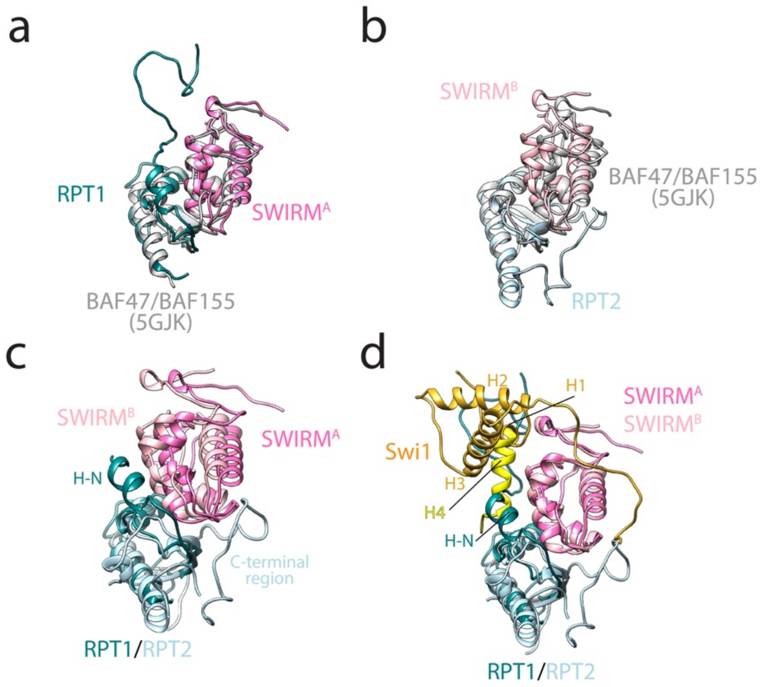


**Extended Data Fig. 7 Structural features of the Arm sub-module of the SWI/SNF complex. a**, The Snf5 RPT1 and Swi3 SWIRM$^{A}$ heterodimer was aligned with the human BAF47/BAF155 crystal structure (PDB ID 5GJK). **b**, The Snf5 RPT2 and Swi3 SWIRM$^{B}$ heterodimer was aligned with the human BAF47/BAF155 crystal structure (PDB ID 5GJK). **c**, RPT1/SWIRM$^{A}$ interface shows slight difference with the RPT2/SWIRM$^{B}$ interface. RPT1 and 2 was aligned, resulting in the SWIRM domains slightly shifting from each other. **d**, Comparing the interfaces between the SWIRM domains and Swi1. The two SWIRM domains was aligned, resulting in Swi1 H4 (yellow; contacting SWIRM$^{B}$) occupying a similar position as Swi1 H1 (gold) and Snf5 H-N on SWIRM$^{A}$. In all panels, structural elements related to RPT1/SWIRM$^{A}$ is depicted using darker colors, whereas RPT2/SWIRM$^{B}$ associated structures is shown in lighter colors.

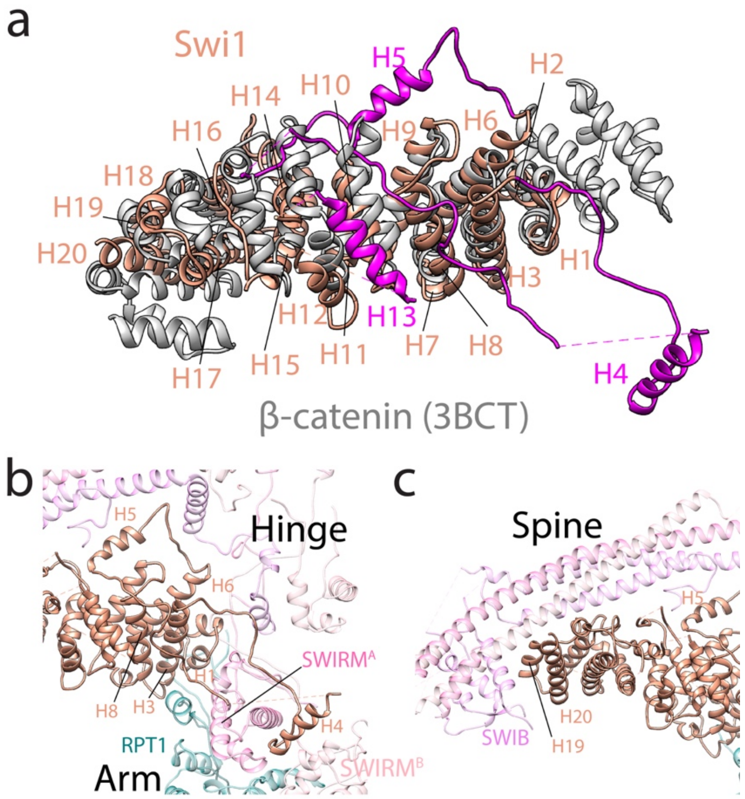


**Extended Data Fig. 8 Structural features of the Core sub-module of the SWI/SNF complex. a**, The Swi1 ARM repeat domain is aligned with β-catenin (gray; PDB ID 3BCT). The insertions of the Swi1 ARM repeat domain are depicted in magenta. **b**, Detailed interaction between the Swi1 ARM repeat domain with the Arm and Hinge sub-modules. **c**, Detailed interaction between the Swi1 ARM repeat domain with the Spine sub-module.

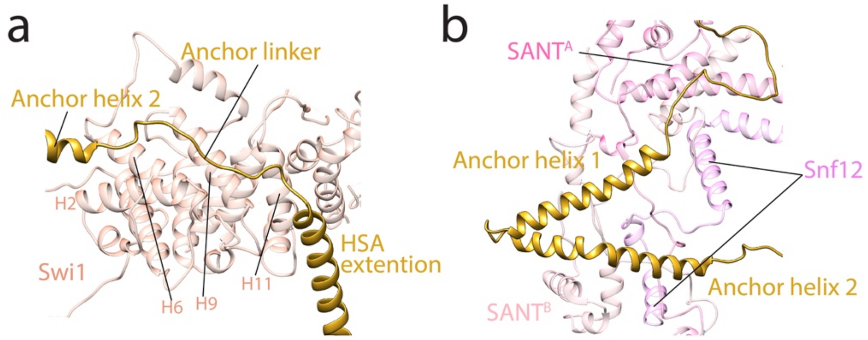


**Extended Data Fig. 9 Interactions between the Snf2 Anchor domain and the rest of the SWI/SNF complex. a**, The Snf2 Anchor linker region interacts with the Swi1 ARM repeat domain. **b**, Snf2 Anchor helices 1 and 2 are sandwiched between the two SANT domains of Swi3 in the Hinge region.

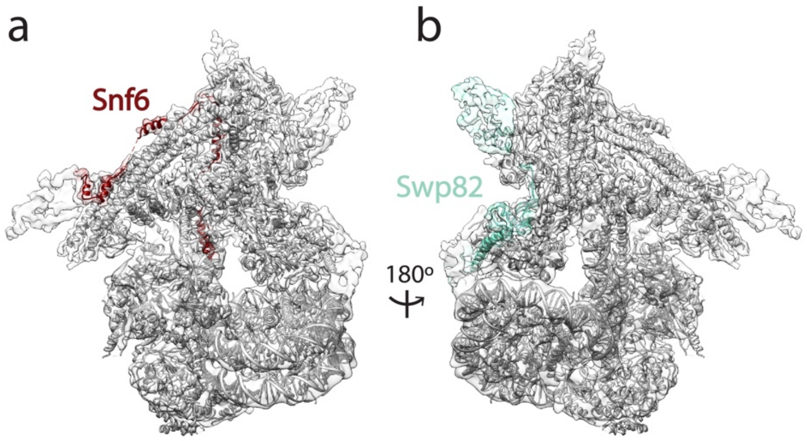


**Extended Data Fig. 10 Yeast specific subunits locate at peripheral locations in the complex.** Snf6 (**a**) and Swp82 (**b**) are positioned at peripheral locations within SWI/SNF. Map and structural models are shown with Snf6 and Swp82 highlighted in **a** and **b**, respectively.

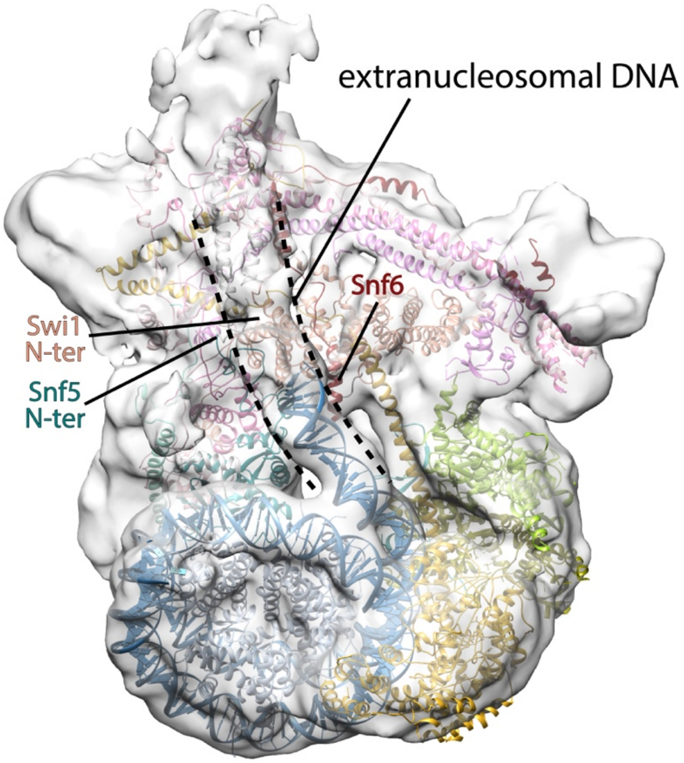


**Extended Data Fig. 11 Density of the extranucleosomal DNA.** The extranucleosomal DNA density is close to Snf6 and is indicated as dashed lines. The N-termini of Swi1 and Snf5 are also labeled. The N-terminal regions of Swi1 and Snf5, which are highly flexible and therefore not resolved in the structure, could take trajectories close to the extranucleosomal DNA.

**Extended Data Table 1. Cα distances of crosslinked residues in SWI/SNF Body module.** Distances are measured in UCSF Chimera. Only those residues that can be mapped onto the SWI/SNF Body structure are measured. For crosslinking pairs involving Swi3, the shortest distances are listed.

**Extended Data Table 2. Data collection, map and model refinement, validation statistics.**

**Extended Data Table 3. Summary of SWI/SNF subunits.**
